# Pnpt1 mediates NLRP3 inflammasome activation by MAVS and metabolic reprogramming in macrophages

**DOI:** 10.1101/2022.05.07.490979

**Authors:** Chia George Hsu, Wenjia Li, Mark Sowden, Camila Lage Chávez, Bradford C. Berk

## Abstract

Polyribonucleotide nucleotidyltransferase 1 (Pnpt1) plays critical roles in mitochondrial homeostasis by controlling mitochondrial RNA (mt-RNA) processing, trafficking and degradation. Pnpt1 deficiency results in mitochondrial dysfunction that triggers a Type I interferon response, suggesting a role in inflammation. However, the role of Pnpt1 in inflammasome activation remains largely unknown. In this study, we generated myeloid-specific Pnpt1-knockout mice, and demonstrated that Pnpt1 depletion enhanced interleukin-1 beta (IL-1β) and interleukin-18 (IL-18) secretion in mouse sepsis models. Using cultured peritoneal and bone marrow-derived macrophages we demonstrated that Pnpt1 regulated NLRP3 inflammasome dependent IL-1β release in response to lipopolysaccharides (LPS), followed by nigericin, ATP or poly (I:C) treatment. Pnpt1 deficiency in macrophages increased glycolysis after LPS, and mt-reactive oxygen species (mt-ROS) after NLRP3 inflammasome activation. Pnpt1 activation of the inflammasome was dependent on both increased glycolysis and expression of the mitochondrial antiviral-signaling protein (MAVS), but not NF-κB signaling. Collectively, these data strengthen the concept that Pnpt1 is an important mediator of inflammation as shown by activation of the NLRP3 inflammasome in mouse sepsis and cultured macrophages.

## Introduction

Mitochondrial homeostasis has gained much attention in studies of immunity (1, 2). Polyribonucleotide nucleotidyltransferase 1 (Pnpt1, previously PNPase or PNPase old-35) is a mitochondria targeted, multi-domain protein containing, 3’-5’ exoribonuclease activity. Pnpt1 localizes in the mitochondrial matrix and in the intermembrane space (3). In the matrix, it takes part in tRNA processing, polyadenylation and degradation; while in the mitochondrial intermembrane space it is involved in the import of different RNA species from the cytoplasm (3-6). Recent studies showed that Pnpt1 mutation or deficiency causes mitochondrial dysfunction that was pathogenic for several diseases (7-9). Patients carrying hypomorphic mutations in the *PNPT1* gene displayed mitochondrial dsRNA (mt-dsRNA) and cytosolic dsRNA accumulation which triggered Type I interferons (IFN) responses in cancer cells and fibroblasts (10, 11).

NLR Family Pyrin Domain Containing 3 (NLRP3) Inflammasomes are multi-molecular signaling complexes involved in the development of atherosclerosis, diabetes and several immune-mediated diseases (12-14). NLRP3 inflammasome activation in macrophages depends on two functionally distinct steps: ‘priming’ and ‘activation’. The ‘priming’ step, classically triggered by microbe-derived LPS, stimulates transcription of proinflammatory cytokines pro-IL-1β and pro-IL-18 and inflammasome component NLRP3 via activation of transcription factor nuclear factor-κB (NF-ĸB). The second step, ‘activation’, is triggered by adenosine triphosphate (ATP), nigericin, monosodium urate and poly I:C, leads to NLRP3 inflammasome assembly followed by caspase-1 activation, and secretion of active pro-inflammatory IL-1β or IL-18 (12, 15).

Mitochondrial dysfunction plays a key role in inflammation by at least three mechanisms. 1) <u>Release of several danger-associated molecular patterns (DAMPs).</u> Under conditions of stress, mitochondria release DAMPs, such as DNA (mt-DNA), double stranded RNA (mt-dsRNA), and reactive oxygen species (mt-ROS), that promote inflammation. (2, 16-19). Production of a mt-ROS signal that contributes to NLRP3 inflammasome activation plays a critical role in the pathogenesis of exaggerated inflammation (20). Highly unstable dsRNA derived from mitochondria damage promotes the production of pro-inflammatory and antiviral cytokines through cytosolic sensors known as retinoic acid-inducible gene I (RIG-I). Cytosolic dsRNA also activates the NLRP3 inflammasome (21, 22), highlighting the important role of Pnpt1, which degrades dsRNA. 2) <u>Function as an essential scaffold for signaling molecules.</u> Mitochondrial antiviral-signaling protein (MAVS) is a critical protein for the production of interferons (23). MAVS acts as a scaffold molecule for NLRP3 oligomerization, which drives caspase-1 activation and IL-1β secretion (17). Depletion of MAVS abrogated Pnpt1-mediated interferon response suggesting that Pnpt1-induced interferon response is also regulated through MAVS signaling (24). Together, these results suggest a potential role of Pnpt1-MAVS axis in the regulation of NLRP3 inflammasome. 3) <u>Control of metabolic reprogramming.</u> NLRP3 inflammasome activation and IL-1β release are regulated by glycolysis and mitochondrial metabolites (2, 25, 26). For example, inhibition of glycolysis by 2-deoxy-d-glucose (2-DG) suppressed LPS-induced IL-1β gene expression and release in macrophages (27). Inflammation is an energy- and redox-intensive process. LPS stimulation induces a glycolytic shift through downregulation of mitochondrial respiration and increasing glycolytic gene expression. This glycolytic shift provides rapid ATP production, and generates nicotinamide adenine dinucleotide phosphate (NADPH) and glutathione (GSH) to protect cells from the production of excess ROS (28, 29). Both NLRP3 and its critical binding partner, NIMA-related kinase 7 (NEK7) are regulated by ROS (30, 31), suggesting that macrophage metabolic reprogramming could regulate NLRP3 inflammasome activation.

Although Pnpt1 depletion results in an interferon response and mitochondrial dysfunction, the role of Pnpt1 in NLRP3 inflammasome activation is unknown. We hypothesized that Pnpt1 is an important mediator of NLRP3 inflammasome activity by regulating mitochondrial function and signaling. To address this hypothesis, we studied inflammation in myeloid cell specific Pnpt1 knockout and wild type mice. These mice were challenged with LPS or LPS+ATP to induce a systemic inflammatory response. The major finding was significantly increased IL-1β production in Pnpt1 myeloid-specific knockout mice compared to wild type mice. Mechanistically, depletion of Pnpt1 in macrophages increased NLRP3 inflammasome activation as measured by IL-1β production through a mechanism requiring MAVS signaling and metabolic reprogramming. These studies describe a novel Pnpt1-dependent pathway in which mitochondrial dysfunction, due to decreased Pnpt1 expression, promotes NLRP3 inflammasome activation.

## Results

### Pnpt1 deletion increased LPS-induced lung inflammation and NLRP3 inflammasome activation

To examine the effects of myeloid cell Pnpt1 depletion, we generated myeloid-specific Pnpt1-knockout mice (Pnpt1^m-/-^) by crossing Pnpt1^flox^/^flox^ mice with LysM-Cre transgenic mice. Whole body Pnpt1 knockout in mice is lethal, but Pnpt1^m-/-^ mice have normal body weight and phenotype. We first used a LPS sepsis mouse model to explore the role of Pnpt1 in lung injury and systemic inflammation (32). Sepsis is a systemic inflammatory response that can cause life-threatening organ dysfunction. Sepsis-related multiple organ dysfunction syndrome, and septic shock significantly increases patient mortality (33). LPS, the major causative agent of gram-negative bacteria-induced sepsis, is capable of stimulating innate immune cells and triggering inflammatory signaling cascades (34). To model sepsis, we injected mice with LPS intraperitoneally (i.p.). After 6 hours, lung tissues and peritoneal lavage fluid were collected to assess the inflammatory response. At baseline, there was no difference in inflammation or lung morphology in wild type mice (WT) compared to Pnpt1^m-/-^ mice (Fig. 1A). In response to LPS, there was significant lung injury as shown by increased pulmonary edema and inflammatory cell numbers (Fig 1A). Macrophage infiltration was significantly increased in Pnpt1^m-/-^ mice after LPS challenge as immunohistochemistry (IHC) showed higher expression of MAC-2 in lung sections of Pnpt1^m-/-^ compared to WT mice (Fig. 1A&B). However, myeloperoxidase (MPO), an abundant ROS generating enzyme, present in neutrophils and macrophages, was not different in Pnpt1^m-/-^ mice compared to WT mice after LPS challenge (Fig. 1A&C).

**Fig. 1.**
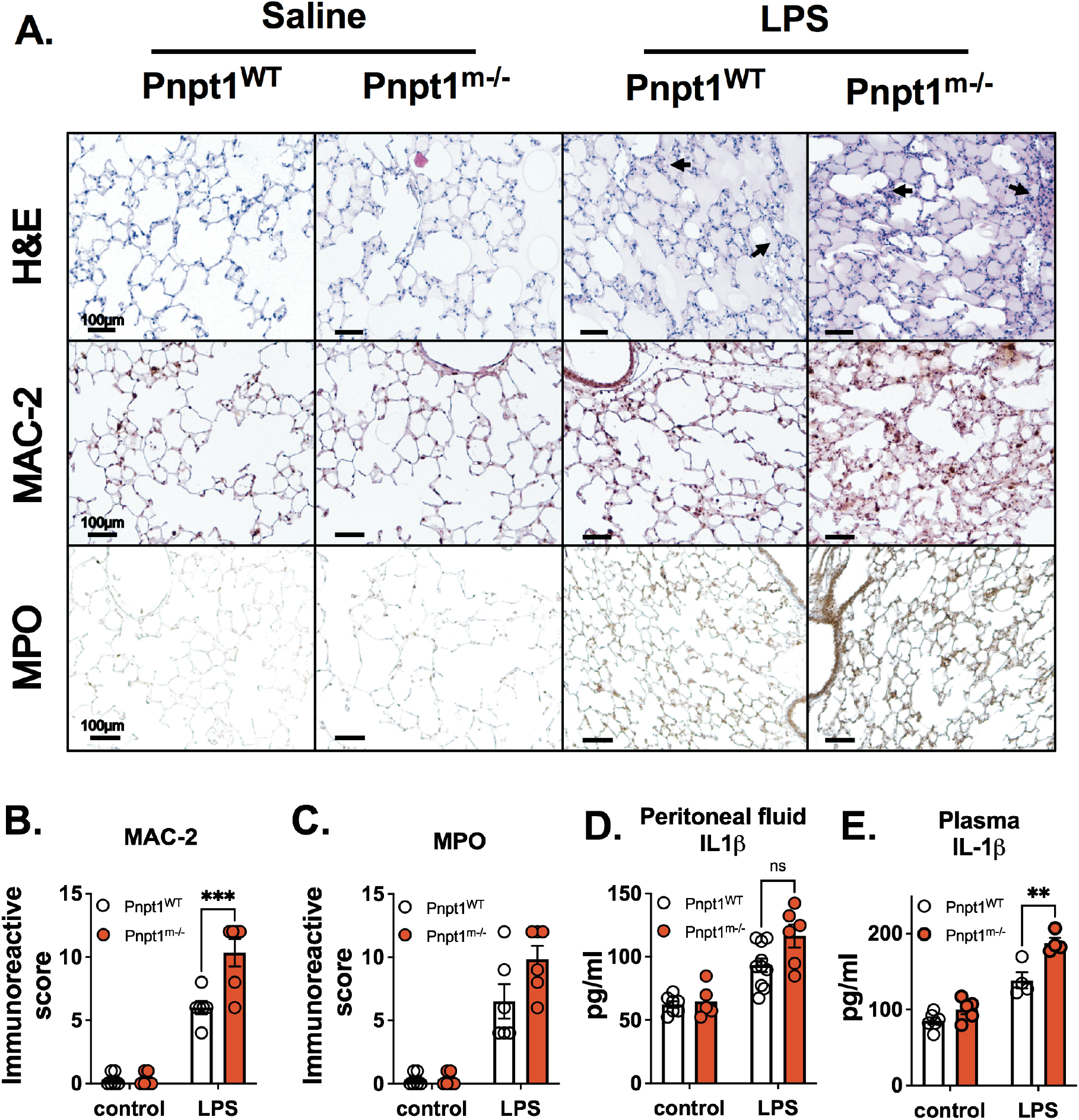
Pnpt1 deletion increased LPS-induced lung and systemic inflammation in vivo. Ten-week-old male Pnpt1^m-/-^ and WT mice were sacrificed 6 h after i.p. injection with 40 mg/kg LPS or saline (control). Whole blood samples were collected and peritoneal cavities were washed with PBS. (A) Representative H&E, Mac-2, and MPO staining of lung sections (B) Quantification of Mac-2 staining (C) Quantification of MPO staining. ELISA of IL-1β level in (D) Peritoneal lavage fluid and in (E) plasma. Statistics in B-E were performed using a 2-way ANOVA and Bonferroni’s post hoc test. N = 6 mice per group. ****P<0.001 between LPS-WT and LPS-Pnpt1^m-/-^ groups. Bars represent mean ± SEM. **p <0.01, ***p <0.001.

To test if Pnpt1 regulated cytokine secretion and systemic inflammation, we measured IL-1β, MCP-1 and IL-6 levels in peritoneal lavage fluid and plasma from Pnpt1^m-/-^ and WT mice, with or without LPS challenge. At baseline, there was no difference between WT and Pnpt1^m-/-^ mice (Fig. 1D–E, Fig. S1A-D). In response to LPS, there were significantly higher levels of IL-1β and MCP-1, but not IL-6 in peritoneal lavage fluid of Pnpt1^m-/-^ compared to WT mice (Fig. 1D, Fig.S1A and B). IL-1β levels in plasma were also higher in Pnpt1^m-/-^ mice compared to WT mice (Fig. 1E), but there was no difference in IL-6 and MCP-1 levels in plasma (Fig. S1C-D).

Intraperitoneal injection of LPS in mice significantly increased production and release of IL-6 and MCP-1. However, the secretion of IL-1β following inflammasome activation requires a second signal (35). We have utilized a clinically relevant, acute *in vivo* sepsis model to study the inflammasome pathway using a combination of LPS and ATP as a second signal (36). To test the possibility that Pnpt1 deletion may increase inflammasome activation *in vivo*, Pnpt1^m-/-^ and WT mice were injected with LPS followed by ATP. Plasma and peritoneal fluid were harvested 1 hr after ATP injection for measurement of IL-1β and IL-18. In response to LPS+ATP, there were significantly increased levels of both cytokines in Pnpt1^m-/-^ vs WT mice (Fig. 2A-D). Collectively, these data show that Pnpt1 deletion enhances inflammasome activation *in vivo* sepsis models.

**Fig. 2.**
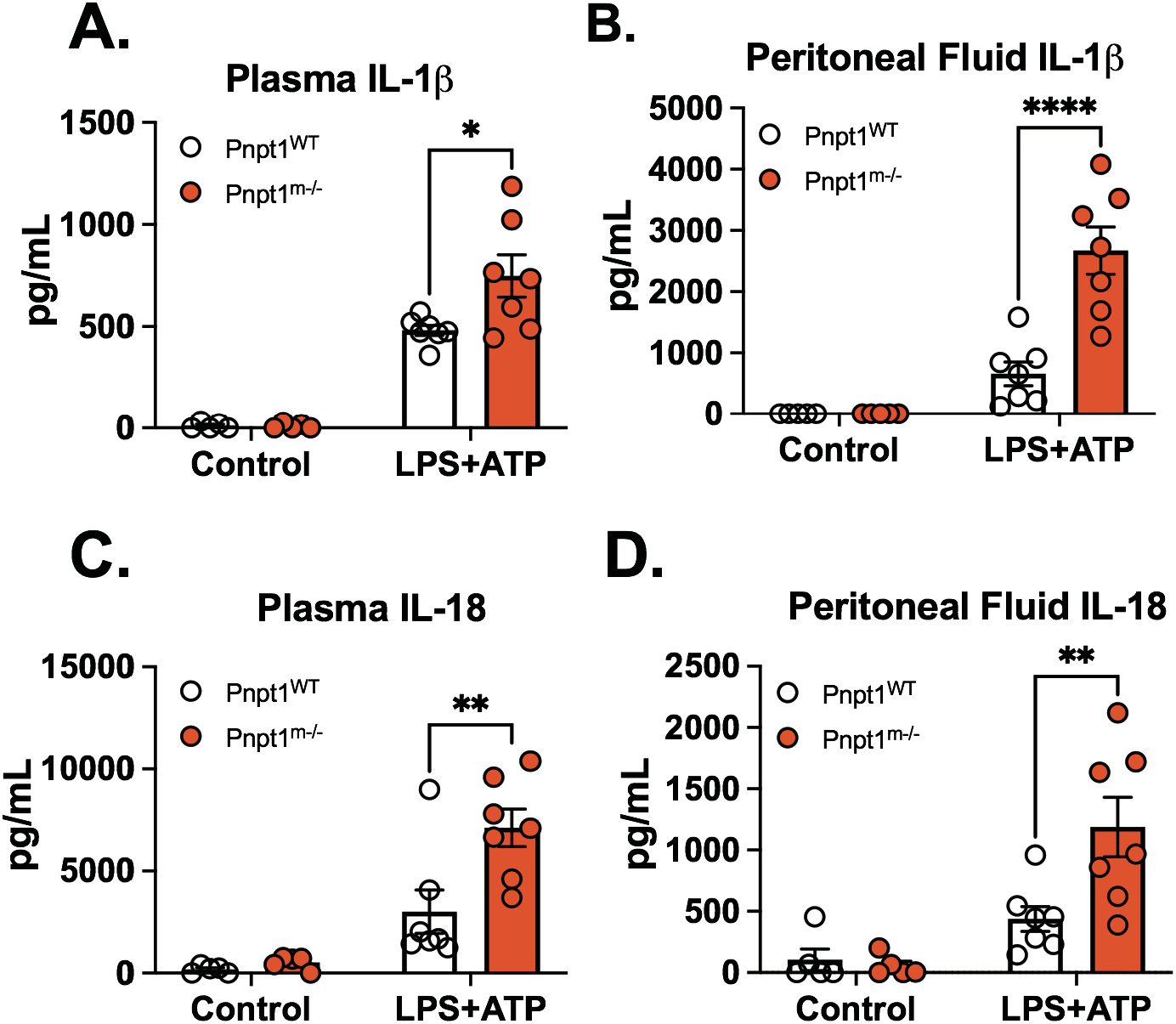
Pnpt1 deletion increased IL-1β and IL-18 production in a mouse sepsis model. Mice were injected with saline (Control) or LPS (10 mg/kg) i.p. for 3 hr followed by ATP (100 mM in 100 μl, pH 7.4) i.p. Plasma and peritoneal fluid were harvested 1 hr after ATP treatment for cytokine assay measured by ELISA. (A) IL-1β in plasma, (B) IL-1β in peritoneal fluid, (C) IL-18 in plasma, (D) IL-18 in peritoneal fluid. (N=5 mice from each control group, and N=7 from each LPS+ATP group). Statistics in A-D were performed using a 2-way ANOVA and Bonferroni’s post hoc test. *P<0.05, **P<0.01, ****P<0.001 between Pnpt1^m-/-^ and WT groups after LPS+ATP treatment. Bars represent mean ± SEM.

### Pnpt1 deficiency increases NLRP3 inflammasome activation independent of NF-κB signaling in macrophages

In our *in vivo* study, we showed Pnpt1 deficiency increased IL-1β in the peritoneal cavity after LPS+ATP injection. NLRP3 inflammasome activation requires a second signal, such as K^+^ efflux (stimulated by nigericin) or ATP that induces NLRP3 oligomerization (12). To study the role of Pnpt1 in NLRP3 inflammasome activation, we used peritoneal macrophages, and mouse bone-marrow-derived macrophages (BMDMs) from WT and Pnpt1^m-/-^ mice, stimulated by LPS and nigericin. IL-1β release was significantly increased in both peritoneal macrophages and BMDMs in Pnpt1^m-/-^ compared to WT mice, (Fig. 3A-B). Similarly, in differentiated human THP-1 macrophages, in which Pnpt1 was depleted using siRNA, there was a two-fold increase in IL-1β release (Fig. 3C). To analyze further the role of Pnpt1, we used ATP or poly (I:C) (a dsRNA mimic) as a second signal to induce inflammasome activation. Both stimuli significantly increased IL-1β secretion in Pnpt1^m-/-^ compared to WT BMDMs (Fig. 3D).

**Fig. 3.**
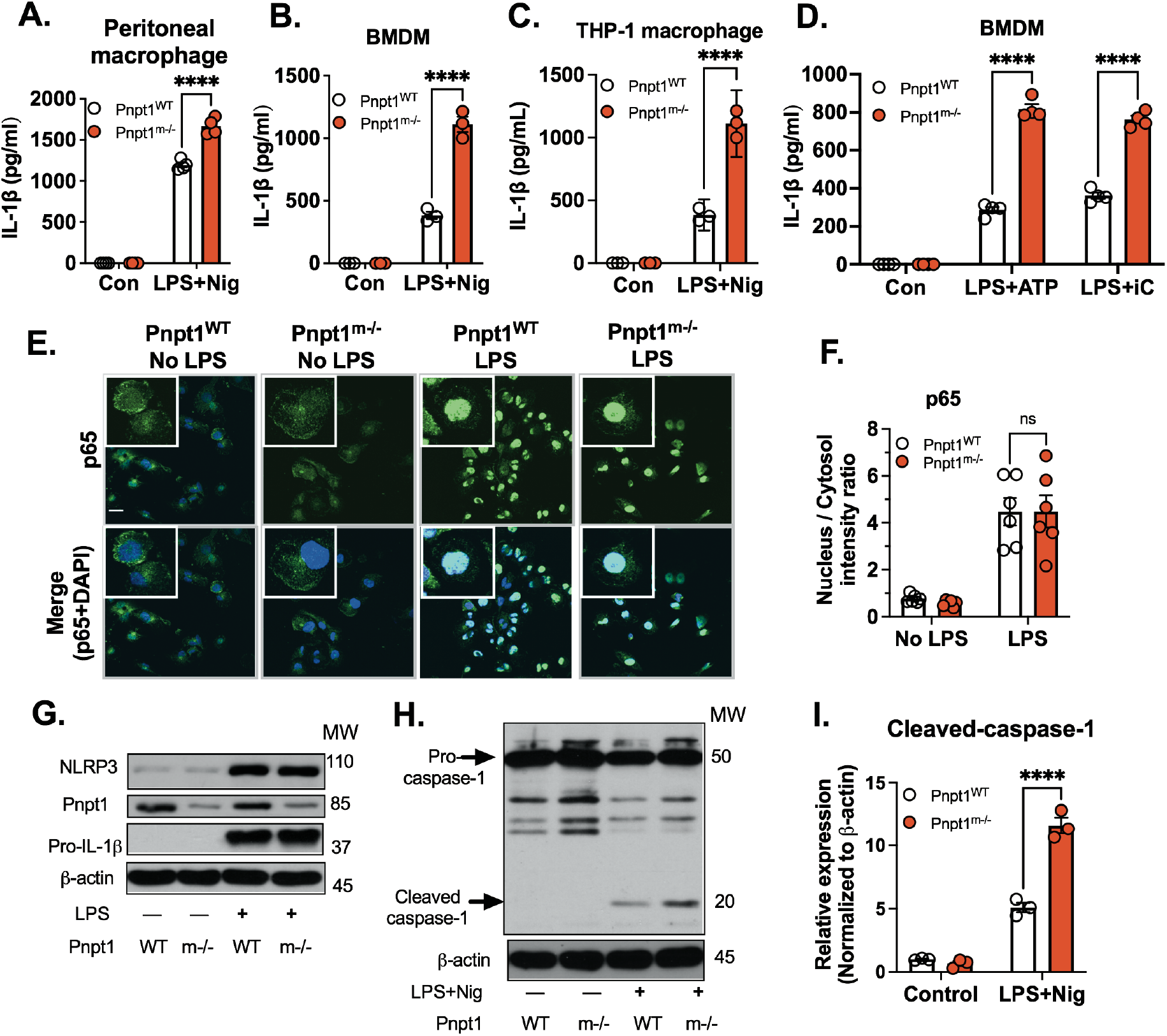
Pnpt1 deletion increased inflammasome activation independent of NF-ĸB pathway. Macrophages from WT and Pnpt1^m-/-^ mice were stimulated with or without LPS (100 ng/mL) for 3 hr and followed by 2 μM nigericin (Nig) for 1 hr. IL-1β release in medium from (A) Peritoneal macrophages, (B) BMDMs, and (C) THP-1 macrophages (Control and Pnpt1 Knockdown). BMDMs from WT and Pnpt1^m-/-^ mice were stimulated with or without LPS (100 ng/mL) for 3 hr. and followed by (D) 2 μM ATP for 1 hr. 10 μM poly (I:C) for 24 hr. IL-1 β release in medium was measured by ELISA. (Con: control). (E) Peritoneal macrophages were stimulated with LPS (100 ng/mL) for 1 hr., NF-κB p65 translocation was measured by immunofluorescence (scale bar: 10 μm), DAPI (blue), and p65 (Green). (F) Quantification using Image J. N=3 experiments. (G) Peritoneal macrophages were stimulated with or without LPS (100 ng/mL) for 3 hr. Protein expression of NLRP3 and pro-IL-1β was analyzed by western blot, and images were representative of three independent experiments. (H) Peritoneal macrophages were stimulated with LPS (100 ng/mL) for 3 hr. followed by 1 μM nigericin (Nig) stimulation for 30 min. Western blots are representative of three independent experiments. Statistics in A-D, F and I were performed using a 2-way ANOVA and Bonferroni’s post hoc test. ****P<0.001 between Pnpt1^m-/-^ and WT groups after stimulation. Bars represent mean ± SEM.

To determine the mechanism of Pnpt1-mediated inflammasome activation, we first focused on NF-κB signaling by assessing p65 nuclear translocation, as well as expression of Pnpt1, NLRP3 and pro-IL-1β in macrophages treated with LPS. In response to LPS (100 ng/mL), there was a 5-fold increase in nuclear translocation of p65, which was not significantly different between Pnpt1^m-/-^ and WT (Fig. 3E-F). In response to LPS, NLRP3 and pro-IL-1β expression were increased (Fig. 3G) in both Pnpt1^m-/-^ and WT macrophages, but there was no increase in Pnpt1 expression (Fig. 3G). Consistent with p65 nuclear translocation, LPS-induced NLRP3 and pro-IL-1β expression were not significantly different between Pnpt1^m-/-^ and WT (Fig. 3G). Inflammasome activation induces the cleavage of pro-caspase-1 to caspase-1. Pnpt1^m-/-^ macrophages showed significantly greater cleavage of caspase-1 than in WT (Fig. 3H-I). Collectively, these findings show that Pnpt1 regulated macrophage inflammasome activation is independent of NF-κB signaling.

### Pnpt1 controls metabolic reprogramming in peritoneal macrophages

The switch from mitochondrial respiration to glycolysis is crucial for the inflammatory response in monocytes, macrophages and dendritic cells (28, 37). Pnpt1 knockout induces metabolic reprogramming by decreasing oxidative phosphorylation and increasing glycolysis (38). However, the effect of Pnpt1 knockout on metabolism in response to a bacterial stimulus, such as LPS, is not clear. To investigate the role of Pnpt1 in metabolic alterations in macrophages, LPS-activated and PBS-treated peritoneal macrophages were assessed using a Seahorse XF-analyzer. Mitochondrial respiration was assessed by recording oxygen consumption rate (OCR, an index of mitochondrial respiration) and extracellular acidification rate (ECAR, an index of glycolysis). Under basal conditions, there was no difference in ECAR between Pnpt1^m-/-^ and WT macrophages (Fig. 4A&B). LPS stimulation significantly increased ECAR in macrophages from both Pnpt1^m-/-^ and WT cells, but the effects were significantly greater in Pnpt1^m-/-^ compared to WT cells (Fig. 4A&B). Furthermore, maximal ECAR was greater in Pnpt1^m-/-^ compared to WT macrophages after LPS stimulation. Consistent with the ECAR data, lactate levels were significantly increased in LPS-treated Pnpt1^m-/-^ macrophages vs WT (Fig. 4D). Although OCR was decreased less in Pnpt1^m-/-^ vs WT (Fig.S2A-B), maximal OCR was similar in Pnpt1^m-/-^ compared to WT macrophages after LPS stimulation (Fig.S2C). As activated macrophages in Pnpt1^m-/-^ have greatly increased glycolysis compared to WT, it is possible that Pnpt1 deletion may alter gene expression related to glucose metabolism. Hypoxia-inducible factor-1 alpha (HIF-1α) and glucose transporter 1 (GLUT1) participate in the regulation of glucose metabolism in LPS-treated macrophages (25, 27, 39). Indeed, following LPS stimulation, HIF-1α and GLUT1 mRNA levels were markedly increased with a significantly greater increase in Pnpt1^m-/-^ compared to WT macrophages (Fig. 4E&F). These findings demonstrate that Pnpt1 regulates metabolic reprogramming in macrophages after LPS stimulation.

**Fig. 4.**
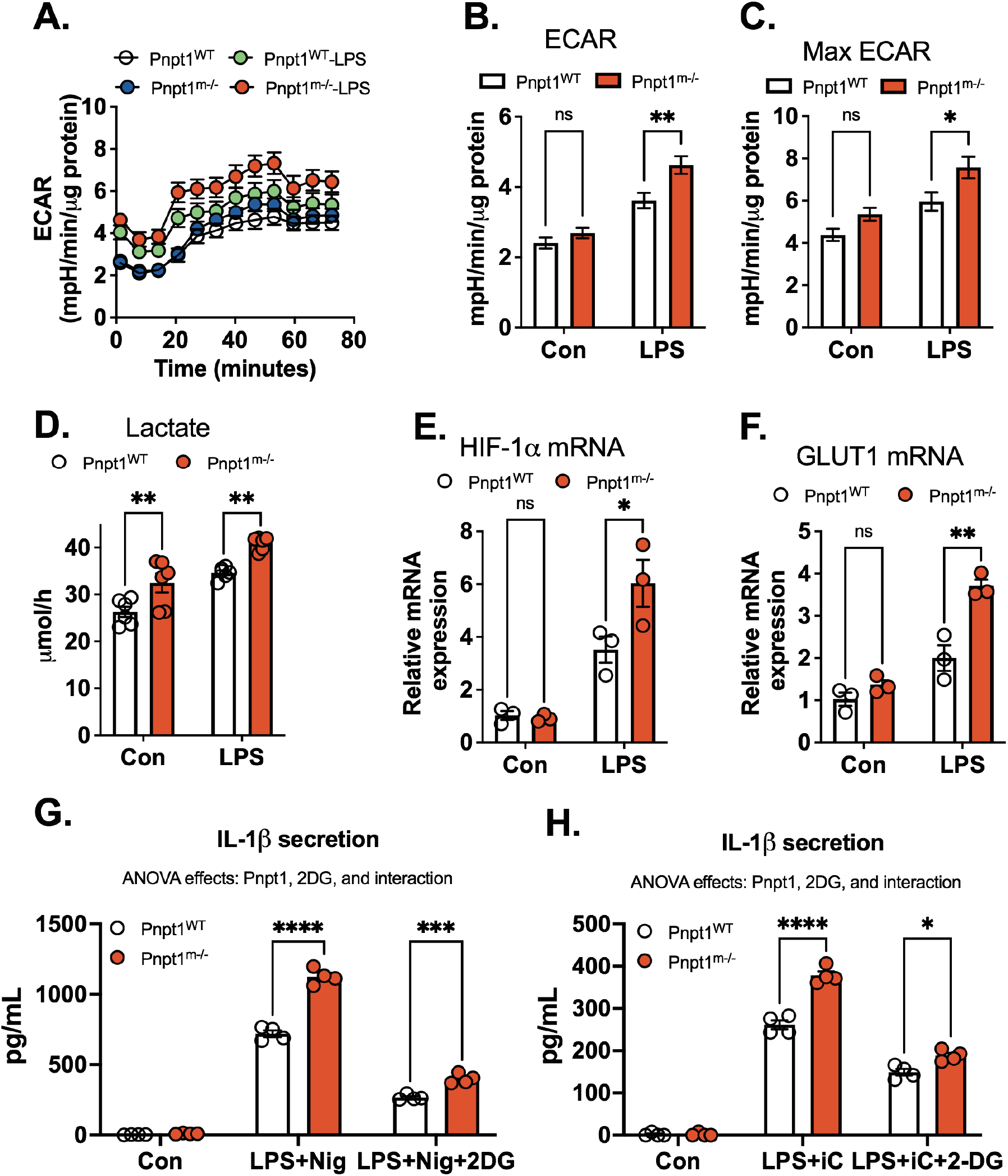
Pnpt1 deletion controls metabolic reprogramming, and contributes to inflammasome activation. Untreated (Con) or LPS (100ng/mL)-activated (3 hr) peritoneal macrophages from Pnpt1^m-/-^ and WT mice were subjected to a mitochondrial stress test using a Seahorse XF-analyzer. (A) A representative Seahorse plot of the mitochondrial stress test assessed by extracellular acidification rate (ECAR), an index of glycolysis after injection of oligomycin (1 μg/ml), carbonyl cyanide-4-(trifluoromethoxy)phenylhydrazone (FCCP, 1 μM), and rotenone (1 μM) plus antimycin (1 μM). (B) Quantification of results from (A) ECAR was defined as the rate at time = 0. (C) Quantification of results from (A) Maximal ECAR was defined as the highest rate at any time. (D) In separate experiments, peritoneal macrophages from Pnpt1^m-/-^ and WT mice were stimulated with LPS for 3 hr. Lactate production in the culture supernatants was measured. (E) The relative gene expression of HIF1α and (F) GLUT1 in response to 100 ng/mL LPS for 3 hr was measured by RT-PCR. (G) Peritoneal macrophages were stimulated with LPS (100 ng/mL) for 3 hr, co-incubated with PBS or 10 mM 2-Deoxy-D-glucose (2DG), then followed by (H) 2 μM nigericin (Nig) for 1 hr, or (I) 10μM poly(I:C) (iC) transfection for 12 hr. IL-1β in the medium was measured by ELISA. Statistics in B-H were performed using a 2-way ANOVA and Bonferroni’s post hoc test. Bars represent mean ± SEM. *p <0.05.

To determine whether Pnpt1-mediated glycolysis contributes to IL-1β release, we used 2-deoxy-D-glucose (2-DG), a competitive inhibitor for hexokinase of the glycolytic pathway (27). Inhibition of glycolysis by 10 mM 2-DG decreased IL-1β release after LPS-nigericin or LPS-poly (I:C) treatment in BMDMs. Statistically there was a significant interaction between Pnpt1 and 2-DG, to inhibit IL-1β release (Fig 4G-H). These results suggest that the increased glycolysis in Pnpt1^m-/-^ after inflammasome activation contributed to IL-1β release in LPS-primed macrophages.

### Pnpt1-medated IL-1β release requires MAVS

Pnpt1 is located in the mitochondrial intermembrane space, regulates the expression of the electron transport chain (ETC) components (6), and Pnpt1 mutations impair mitochondrial function (8, 9). To determine the role of Pnpt1-mediated mt-ROS generation in inflammasome activation, we measured mitochondrial superoxide (MitoSOX Red™ Indicator) and hydrogen peroxide (Amplex® Red Hydrogen Peroxide/Peroxidase Assay). Under baseline conditions, there was no difference in mt-ROS between Pnpt1^m-/-^ and WT peritoneal macrophages (Fig. 5A&B). However, in response to inflammasome activation (LPS+Nig), there was a ∼4-fold increase in mt-ROS, which was significantly greater in Pnpt1^m-/-^ compared to WT cells (Fig. 5A&B). Hydrogen peroxide production showed similar results (Fig. 5C). Mitochondrial membrane potential (ΔΨm), analyzed by JC-1 fluorescence, was similar at baseline (Fig. 5D), and decreased following LPS+Nig treatment. The decrease was significantly greater in Pnpt1^m-/-^ compared to WT cells (Fig. 5D). MAVS oligomerizes in response to mt-ROS, and oligomerization facilitates the recruitment of NLRP3 inflammasome components to the mitochondria, enhancing inflammasome formation and activity (17, 40, 41). Because MAVS activation is required for the IFN response, we determined the role of MAVS in interferon activation in the context of Pnpt1 deficiency. Pnpt1 knockdown in 293T cells significantly increased mRNA levels of IFN-β and interferon-induced gene 15 (ISG-15) compared to control group (Fig. S3A-B). siRNA knockdown of MAVS abrogated the interferon response in Pnpt1 deficient cells (Fig. S3A-B). These data suggest that MAVS is an essential effector of the Pnpt1-mediated IFN response. Next, we determined whether Pnpt1-mediated IL-1β release requires MAVS. Pnpt1 or MAVS expression in THP-1 macrophages was depleted with siRNA followed by stimulation with LPS +Nig. We showed that reduced MAVS expression decreased inflammasome activation as measured by IL-1β release in Pnpt1 depleted cells (Fig 5E). Consistent with these results, MAVS knockdown decreased Pnpt1-mediated IL-1β release after poly (I:C) treatment (Fig 5F). To determine if Pnpt1 deficiency triggers the IFN response in macrophages, we measured mRNA levels of IFN-β and ISG-15 after cytosolic poly (I:C) treatment in LPS primed macrophages. We found that under baseline conditions, there was no difference in IFN-β and ISG-15 gene expression in Pnpt1^m-/-^ compared to WT BMDMs (Fig. S3C-D). However, in response to cytosolic poly (I:C), IFN responses were significantly greater in Pnpt1^m-/-^ compared to WT cells (Fig. S3C-D). Furthermore, MAVS depletion decreased Pnpt1-mediated ISG-15 gene expression in THP-1 macrophages after cytosolic poly (I:C) treatment (Fig 5G). Collectively, these results suggest that MAVS expression is required for Pnpt1-mediated NLRP3 inflammasome activation (IL-1β release) and the IFN response (IFN-β And ISG-15 expression).

**Fig. 5.**
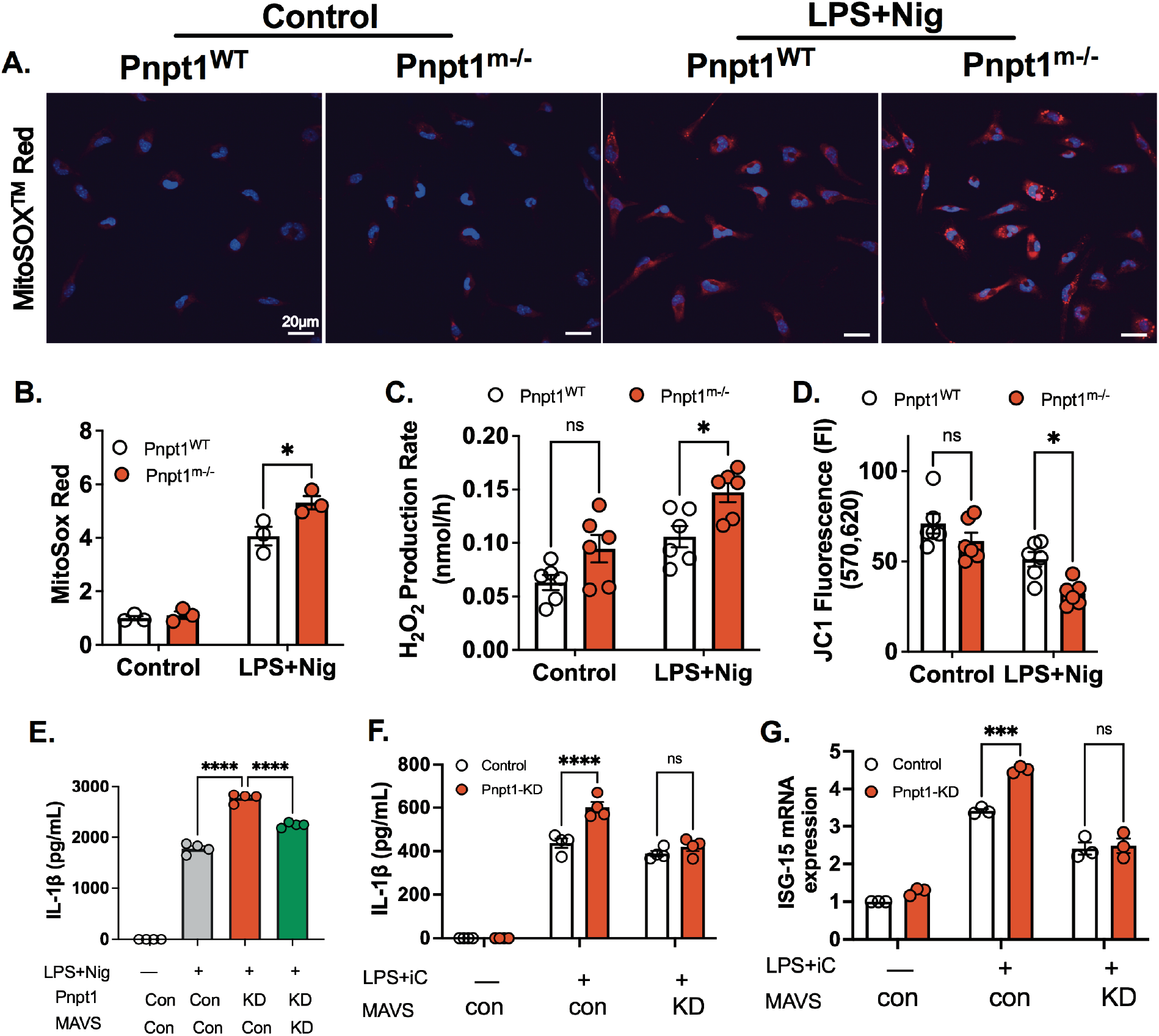
Pnpt1 deficiency causes mitochondrial dysfunction, ROS generation, MAVS activation, and increases IL-1b secretion. Peritoneal macrophages from Pnpt1^m-/-^ and WT mice were primed with 100 ng/ml LPS for 3 h followed by nigericin (2 μM) for 1 hour. (A) Fluorescence microscopy analysis for mtROS production. Nuclei were stained with DAPI (blue) and mitoSOX (red). Scale bar, 20 μm. (B) Quantification of mitoSOX Red fluorescence intensity normalize to control. (C) Hydrogen peroxide production was measured in the culture supernatants of untreated (Control) or LPS+Nig (Nig) activated peritoneal macrophages from Pnpt1^m-/-^ and WT mice. (n=3 experiments with duplicate samples from each group). (D) Quantitative analysis of mitochondrial membrane potential by JC1 fluorescence in peritoneal macrophages (n=3 experiments with duplicate samples from each group). (E) THP-1 macrophages were treated with scrambled, Pnpt1 or MAVS siRNA for 48 hr., and stimulated with LPS (100 ng/mL) for 3 hr. followed by (E) nigericin (Nig, 2 μM) for 1 hr or (F) 10μM poly(I:C) transfection (iC) for 12 hr. The amount of IL-1β in the medium was measured. (G) The amount of ISG mRNA from cell lysates after stimulation with LPS plus poly (I:C) was measured in cells treated with scrambled or MAVS siRNA (n=4 experiments, control: con; knockdown: KD). Statistics in B-D and F-G were performed using a 2-way ANOVA and Bonferroni’s post hoc test. Statistics in E were performed using a one-way ANOVA and Bonferroni’s post hoc test. N=4 experiments. Bars represent mean ± SEM. *p <0.05, ***p <0.001, ****p <0.001.

## Discussion

The major finding of this study is that Pnpt1 deficiency enhances NLRP3 inflammasome activation in cultured macrophages and mouse sepsis models (Fig. 6). To study the role of Pnpt1 in inflammasome signaling, we generated a myeloid cell lineage specific deletion of Pnpt1 in mice (Pnpt1^m-/-^). We showed that Pnpt1^m-/-^ mice injected with LPS or LPS+ATP had excessive lung inflammation, as well as increased IL-1β release in plasma and peritoneal fluid compared to WT mice. Using cultured peritoneal macrophages and BMDMs, we showed that Pnpt1 deletion increased both mitochondrial dysfunction after LPS+nigericin treatment and LPS-induced glycolysis. Furthermore, inhibiting glycolysis using 2-DG, or depletion of MAVS, significantly decreased the Pnpt1 effect on IL-1β release after nigericin or cytosolic poly(I:C) treatment in LPS-primed macrophages. These results show that Pnpt1 is a novel mediator of the mitochondria-immune response. Specifically, we found that Pnpt1 acts as a negative regulator of the NLRP3 inflammasome in a mouse sepsis model and in cultured macrophages by controlling the MAVS pathway and glucose metabolism (Fig. 6).

**Fig. 6.**
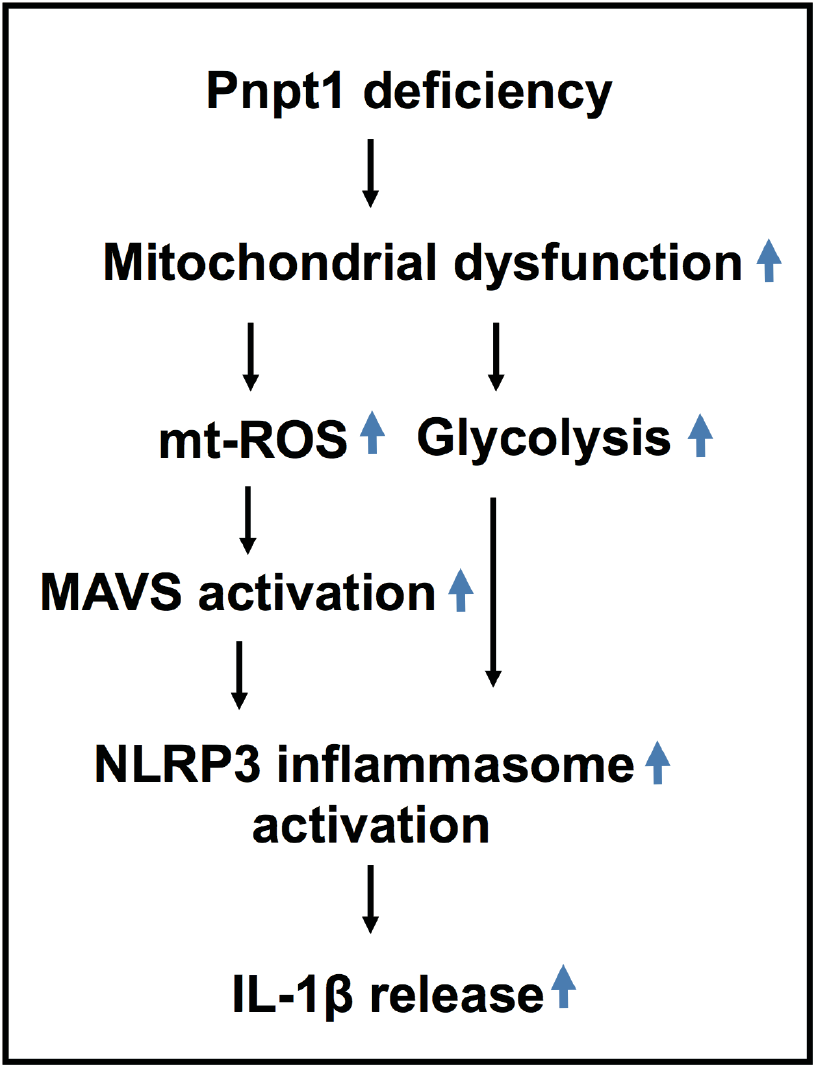
Model of Pnpt1-mediated NLRP3 inflammasome activation by MAVS and glycolysis. After stimulation with LPS and either nigericin or poly(I:C), NLRP3 inflammasome activation and IL-1β release were significantly upregulated in Pnpt1 deficient macrophages by increased MAVS activation and glycolysis.

A novel finding was the requirement for MAVS to stimulate assembly of the NLRP3 inflammasome in the setting of decreased Pnpt1. Our data is consistent with previous reports that decreased Pnpt1 stimulated the MAVS-dependent IFN response (10). MAVS oligomerizes on the outer mitochondrial membrane to create a scaffold for assembly of the NLRP3 inflammasome (17). However, the role of the Pnpt1-MAVS pathway in regulation of inflammasome activation was unknown. In this study, MAVS knockdown decreased Pnpt1-mediated IL-1β release after nigericin or poly (I:C)-induced inflammasome activation suggesting a Pnpt1-MAVS axis that regulates NLRP3 inflammasome activity. There may be other connections between Pnpt1 and MAVS, such as mt-dsRNA, which may be released due to increased mitochondrial membrane permeability in the setting of Pnpt1 depletion (10, 42). This could cause accumulation of cytosolic dsRNA triggering a MDA-5 and MAVS dependent NLRP3 inflammasome response.

MAVS can also be activated by dsRNA-independent mechanisms. For example, generation of mt-ROS can trigger MAVS oligomerization and mediate an inflammasome response (40, 43). This mechanism was supported by findings that antioxidants can suppress-ROS-induced MAVS oligomerization and inflammasome activation (40). In our study, mt-ROS and mitochondrial membrane potential were further enhanced in Pnpt1 deficient cells compared to WT cells after inflammasome activation. It is possible that MAVS senses mt-ROS, which in turn promotes NLRP3 inflammasome activation.

Glycolysis is increased during inflammatory activation of immune cells, including macrophages (1, 39, 44). LPS stimulation results in rapid transcriptional and post-translational responses that involve genes regulating the metabolic shift from oxidative phosphorylation to glycolysis (39). In this study, we showed that Pnpt1 deficiency increased LPS-induced lactate production and glycolysis, as well as HIF-1α gene expression and its transcriptional activity, as measured by GLUT1 expression. Our observations are consistent with reports that Pnpt1 deficiency caused a glycolytic shift in stem cells (45) and fibroblasts (38). In our study, inhibition of glycolysis by 2-DG decreased IL-1β release after LPS+nigericin or LPS+poly (I:C) treatment of BMDMs, suggesting that the hyperinflammatory phenotype in Pnpt1 deficient macrophages may involve metabolic reprogramming. HIF-1α signaling and glycolytic activity play critical roles in both viral and innate immune responses. For example, HIF-1α is important for the full expression of many glycolytic genes including GLUT1, hexokinase, and GAPDH. The increase in glucose uptake and the accumulation of succinate enhance the production of IL-1β (27). It is thus likely that the increased IL-1β release following inflammasome activation in Pnpt1 deficient macrophages depends on a combination of glycolysis and mitochondrial signaling.

Pnpt1 depletion in HeLa cells leads to accumulation of dsRNAs both in mitochondria and the cytoplasm that triggers a type I IFN response through MAVS signaling (10). In contrast to the previous study, we found that Pnpt1 deletion alone in macrophages did not induce an IFN response. Furthermore, Pnpt1 deletion in peritoneal macrophages did not affect the NF-ĸB pathway (as measured by LPS-induced p65 nuclear translocation), or NLRP3 and pro-IL-1β protein expression. These results suggest that NF-ĸB was required for the expression of inflammasome components, but was not important for the observed Pnpt1-mediated effects. LPS treatment triggers IL-6 and MCP-1 secretion, but mature IL-1β production is a multi-step process that is dependent on NLRP3 inflammasome activation. In our mouse sepsis models, IL-1β levels in plasma and peritoneal fluid were higher in Pnpt1^m-/-^ mice compared to WT mice (Fig. 1E), but there was no difference in IL-6 and MCP-1 levels supporting a mechanism whereby Pnpt1 regulates NLRP3 inflammasome activation independent of the NF-ĸB pathway.

Pnpt1 is evolutionarily conserved and has several well described mitochondrial functions (46). Increased mt-ROS and increased OCR and ECAR were expected in Pnpt1 depleted cells. The magnitude of change in baseline was small, but the effect of Pnpt1 deletion was obvious after LPS alone or LPS+ATP. These effects were demonstrated both in cultured macrophages and an animal model of sepsis. Because expression of Pnpt1 can be found in many different cells, this pathway is likely to be important not only in macrophages, but also in most cells activated by DAMPs that stimulate the inflammatory response. Future studies will be required to characterize the specific role of Pnpt1 in other cell types involved in inflammation and the pathogenesis of disease.

Collectively, our results indicate that Pnpt1 is involved in the regulation of NLRP3 inflammasome activation by MAVS and a glycolytic shift, and strengthens the significance of mitochondrial signaling and glucose metabolism in the control of inflammation. Further understanding the interplay between Pnpt1, mitochondrial homeostasis, and immune responses is likely to yield new insights into the pathogenesis of inflammatory diseases.

## Methods

### Mice

The myeloid-specific Pnpt1-knockout mice (Pnpt1^m-/-^) were generated by crossing Pnpt1^flox^/^flox^ mice with LysM-Cre transgenic mice. All mice were in C3H/HeJ background and maintained under specific pathogen–free conditions with free access to pellet food and water, and kept on a 12 hr light–dark cycle. Ten-week-old male Pnpt1^m-/-^ mice and Pnpt1^WT^ littermates were used. Animal welfare and experimental procedures were carried out in accordance with the Guide for the Care and Use of Laboratory Animals (National Institutes of Health, USA) and ethical regulations of University of Rochester.

### Chemicals and reagents

LPS (Escherichia coli O127:B8, L3129), Nigericin (n7143), PMA (Phorbol 12-myristate 13-acetate) (P8139), ATP (A7699), Poly(I:C) (P1530-25MG), oligomycin (O4876), carbonyl cyanide-4-(trifluoromethoxy)phenylhydrazone (FCCP, C2920), rotenone (R8875), antimycin (A8674), and 2-Deoxy-D-glucose (D6134-5G) were purchased from Sigma-Aldrich (St. Louis, MO). Lipofectamine 2000 (11668027) were purchased from Thermo Fisher Scientific (Waltham, MA). Enzyme-linked immunosorbent assay (ELISA) kits for mouse IL-6 (431304), mouse MCP-1(432704), mouse IL-1β (432604) and human IL-1β (437004) were purchased from BioLegend. Mouse IL-1β (BMS6002TWO) and mouse IL-18 (88-50618-22) ELISA Kits were from Thermo Fisher Scientific. Anti-mouse/human Mac-2 (Galectin-3) (CL8942AP, 1: 200 dilution) was purchased from Cedarlane Laboratories. Lab Vision™ Myeloperoxidase (MPO), Rabbit Polyclonal Antibody (RB-373-A, 1:300 dilution) was bought from Thermo Fisher Scientific. Anti-Pnpt1 (14487-1-AP) was purchased from Proteintech (Rosemont, IL). Anti-NLRP3 (AG-20B-0014-C100, 1 : 1,000 dilution), and anti-Caspase-1 (p20) (mouse) (AG-20B-0042-C100, 1: 1,000 dilution) were purchased from AdipoGen. NF-ĸB p65 (D14E12) XP® Rabbit mAb (8242S, 1: 1,000 dilution), β-actin (4970, 1:2000 dilution), Anti-mouse IgG, HRP-linked Antibody (7076), Anti-rabbit IgG, HRP-linked Antibody (7074), were purchased from Cell Signaling Technology (Danvers, MA). IL1-β antibody (GTX10750, 1: 1,000 dilution for immunoblotting) was purchased from GeneTex (Irvine, CA). Anti-goat IgG, HRP-linked Antibody was purchased (805-035-180) from Jackson Immuno Research (West Grove, PA), DAPI-fluoromount-G (0100-20) was purchased from Southern Biotech (Birmingham, AL).

### Mouse sepsis model

#### LPS-induced sepsis

Sepsis was induced in Pnpt1^WT^ and Pnpt1^m–/–^ mice by intraperitoneal injection with 40 mg/kg bacterial endotoxin (E. coli O127:B8, LPS) 6 h before harvesting.

#### LPS/ATP-induced sepsis

Sepsis was induced in mice by intraperitoneal (IP) injection with LPS (10 mg/kg) for 3 hr. followed by 100 mM/100μl per mouse, pH 7.4 ATP (Sigma-Aldrich, A7699) IP injection as previously described (36). Whole blood samples were collected in tubes with 10 μL 500 mM EDTA and centrifuged at 1000 g for 10 min at 4°C. Plasma was collected and aliquoted for cytokine assay. Peritoneal fluids were harvested by washing mouse peritoneal cavity with 2mL ice-cold PBS supplemented with 1mM EDTA. The cell suspension was centrifuged at 500 g for 5 min at 4°C, and the supernatant was collected and aliquoted for cytokine assay.

### Histopathological analysis of lung tissues

Lung tissues of mice from each group were harvested aseptically 6 h after LPS/PBS injection and fixed in 10% formalin overnight. Fixed tissues were embedded in paraffin and then sliced into 5 μm-thick sections. Sections were dealt with hematoxylin and eosin (H&E) staining or immunohistochemical (IHC) staining according to standard protocol. H&E and IHC images were observed under light microscopy (Nikon ECLIPSE 80i). IHC staining was independently assessed by three experienced investigators. Immunoreactive score (IRS) was determined by multiplying the staining intensity in four gradations with the percentage of positive cells. Thus, a score from 0 to 12 points was obtained (47).

### Cell culture

#### Peritoneal macrophage isolation

Peritoneal macrophages were stimulated by injecting mice intraperitoneally with 1 mL of sterile 2% Bio-Gel® P Polyacrylamide Beads (Bio-Rad, 150-4174, Hercules, CA), and were subsequently harvested 4 days later by flushing the peritoneal cavity with 5 ml ice-cold phosphate-buffered saline (PBS) and plated in 6-well plates (5 × 10^5^ cells/well) in RPMI medium 1640 (Thermo Fisher Scientific, 11875-093, Waltham, MA) supplemented with 10% fetal bovine serum (Gibco, 10437-028, Waltham, MA), 100 U/mL penicillin, and 100 μg/mL streptomycin, 1% pyruvate (Thermo Fisher Scientific, 11360070). The next day, cells were washed with serum-free medium twice before use (48).

#### Bone marrow progenitor cell isolation and bone marrow-derived macrophage (BMDM) differentiation

BMDMs preparation was performed as previously described (49). L929 conditioned media which contains the macrophage growth factor M-CSF, was prepared by culturing L929 cells (ATCC) in complete DMEM (Thermo Fisher Scientific, MT10013CV) supplemented with 10% FBS, and 1% penicillin and streptomycin for 10 days at 37°C, 5% CO2. The L929 conditioned media, was collected, filtered (Vacuum Filter/Storage Bottle System, Corning, 431153, Corning, NY), and stored at −80 °C until required. For isolation of BMDMs, tibias and femurs were removed from both male and female mice and flushed with media using a 26-gauge needle. Bone marrow was collected at 500 x g for 2 min at 4 °C, resuspended with complete DMEM medium and filtered through a 70-μm cell strainer (VWR international, 10199-657, Radnor, PA). Bone marrow progenitor cells were cultured in 100 mm dishes for 6-7 days in 70% complete DMEM medium and 30% L929-conditioned medium. Fresh medium (5 mL) was added on day 3. BMDMs were collected by scraping in cold EDTA (1 mM). After centrifugation, BMDMs were seeded into 12-well plates at a density of 1.6 × 10^5^ cells/well in DMEM and incubated overnight before use.

#### THP-1 differentiated macrophage

Human THP-1 monocytes were differentiated into macrophages by 24 hr. incubation with 100 nM PMA (Sigma-Aldrich) in complete RPMI medium at 1.6 × 10^5^ cells/well in 12-well plates. Cells were washed twice with 1x PBS and incubated with complete RPMI medium without PMA for 24 hr prior to any experiment.

### LPS stimulation

Macrophage cultures were rinsed twice with medium without serum and antibiotics (RPMI 1640 for peritoneal macrophages, THP-1 macrophages, and DMEM medium for BMDMs). Cells were exposed to LPS (100 ng/mL) for 3 hr in appropriate growth media without serum and antibiotics.

### Inflammasome stimulation

To activate the NLRP3 inflammasome, cells were primed for 3 h with LPS (100 ng/mL) followed by addition of 2 μM nigericin or 2 mM ATP for 30-60 min. To activate the NLRP3 inflammasome by cytosolic poly(I:C), macrophages were stimulated with LPS (100 ng/mL) for 3 hr followed by intracellular poly(I:C) delivery (10 μg/mL) by transfection using lipofectamine 2000 (Invitrogen, 11668027), according to the manufacturer’s protocol, in OptiMEM (Gibco, 31985-070) for 12-24hr.

### Western blot

Cell lysates were prepared using Cell Lysis Buffer (Cell Signaling Technology, 9803S) containing protease and phosphatase inhibitors (Sigma Aldrich, St. Louis, MO, USA), followed by centrifugation at 12,000 rpm for 10 min. Protein concentrations were measured by BCA protein assay (Pierce™ BCA Protein Assay Kit, Thermo Fisher Scientific, 23225). Cell lysates were separated on 10% SDS-PAGE and were transferred onto nitrocellulose transfer membranes. After blocking using 5% BSA, the membranes were incubated with primary antibodies at 4 °C overnight. Reactive proteins were detected using appropriate secondary antibodies followed by chemiluminescent detection and visualization. The protein signals were quantified by ImageJ software.

### Analysis of mRNA expression

RNA isolation was performed using an RNeasy mini kit (Qiagen GmbH, Hilden, Germany). A total of 1 μg RNA was reverse transcribed using iScript™ cDNA Synthesis Kit (Bio-Rad, 1708891, Hercules, CA, USA). Quantitative PCR amplification reactions contained a target specific fraction of cDNA and 1 μM forward and reverse primers in iQ™ SYBR® Green Supermix (Bio-Rad, 1708882), and analyzed using real-time qPCR. β-actin was used as control for normalization. The ΔΔCT method was used to quantify mRNA levels. The primers used are as follows: Mouse β-actin (Forward: TTCAACACCCCAGCCATGT, Reverse: GTAGATGGGCACAGTGTGGGT); Mouse GLUT1(Forward:GGTTGTGCCATACTCATGACC, Reverse: CAGATAGGACATCCAGGGTAGC); Mouse HIF-1α (Forward: CCCATTCCTCATCCGTCA AATA, Reverse: GGCTCATAACCCATCAACTCA); Mouse ISG15 (Forward: CATCCTGGTGAGGAACGAAAGG, Reverse: CTCAGCCAGAACTGGTCTTCGT); Mouse IFN-β (Forward: CTCAGCCAGAACTGGTCTTCGT, Reverse: CTCAGCCAGAACTGGTCTTCGT); Human ISG15 (Forward: CTCAGCCAGAACTGGTCTTCGT, Reverse: CTCAGCCAGAACTGGTCTTCGT); Human IFN-β1 (Forward: ATGACCAACAAGTGTCTCCTCC, Reverse: GCTCATGGAAAGAGCTGTAGTG); Human β-actin (Forward: TGTCCCCCAACTTGAGATGT, Reverse: TGTGCACTTTTATTCAACTGGTC);

### RNA interference

Differentiated THP-1 monocytes were treated 1 μL/mL lipofectamine RNAiMax (Invitrogen,13778-075) and 3 nM siRNAs in OptiMEM (Gibco, 31985-070). Human siPnpt1 (HSS131759), Human Silencer® Select MAVS (s33179), control negative siRNA (4390844) were purchased from Thermo Fisher Scientific.

### Metabolism analysis

Glycolytic and mitochondrial activity of macrophages were measured using a Seahorse extracellular flux 96 analyzer (Seahorse Bioscience). Peritoneal macrophages were seeded at a density of 1 ×10^5^ cells/ well in 100 μl RPMI 1640 medium, and allowed to adhere for 2 hours. Following washing with PBS, 200 μl RPMI 1640 was added and cells incubated overnight. The next day, cells were washed twice and filled with Seahorse XF base medium (Agilent, 103334-100) (Santa Clara, CA, USA) containing 2g/L glucose, 1X glutamax (Thermo Fisher Scientific, 1798324) and 1X sodium pyruvate (Thermo Fisher Scientific, 11360070) with or without LPS for 3 hours in CO2 free incubator. Oxygen consumption rate (OCR) and extracellular acidification rate (ECAR) were measured following the injection of oligomycin (1 μg/ml), carbonyl cyanide-4- (trifluoromethoxy) phenylhydrazone (FCCP, 1 μM), and rotenone (1 μM) plus antimycin (1 μM).

### Lactate measurement

Mouse peritoneal macrophages were seeded in six-well plates (5 × 10^5^ cells/well) and treated with or without LPS for 3 hr. Lactate dehydrogenase (LDH) in the supernatant was measured using L-Lactate Assay Kit (1200011002, Eton Bioscience, San Diego, CA, USA) according to the manufacturer’s instructions.

### Detection of mitochondrial ROS

A total of 2.5×10^5^ peritoneal macrophages/well were seeded in a 35 mm diameter cover glass dish (Mattek Corp. Ashland, MA). After 24 h of pre-incubation, cells were treated with 100 ng/ml LPS for 3 hr. Culture medium was exchanged for Hank’s balanced salt solution with calcium and magnesium (HBSS/Ca/Mg), containing 5 μM MitoSOX Red™ Mitochondrial Superoxide Indicator (Molecular Probes; Thermo Fisher Scientific M36008, Inc., Waltham, MA, USA), and the cells were incubated for 10 min at 37°C. Following three washes, cells were stimulated with nigericin (2 μM) for 30 min. Cells were fixed in 4% paraformaldehyde for 10 min. After three washes with 1x PBS, cells were mounted using fluoromount-G-DAPI. Cellular fluorescence (510/580 nm) intensity of each of 10 randomly selected cells was detected using a confocal inverted fluorescence microscope (Olympus Corporation, Tokyo, Japan) under ambient temperature. Fluorescence intensity profiles were obtained using the plot profile plugin of ImageJ software. Data were normalized against maximum intensity values in each channel. Results represent pooled data from three independent experiments and data were analyzed independently by two investigators to assure accuracy in the quantification process.

### Determination of extracellular hydrogen peroxide production

Extracellular hydrogen peroxide was quantified using an Amplex® Red Hydrogen Peroxide/Peroxidase Assay Kit (Thermo Fisher Scientific, A22188), according to the manufacturer’s instructions. Briefly, macrophages were seeded in 6-well plates (5 × 10^5^ cells/well), activated by the addition of LPS for 3 hr and supernatants collected. Standard curve samples, controls, and experimental samples (50 μl of each) were pipetted into individual wells of a microplate, along with 100 μM Amplex® Red reagent and 0.2 U/mL HRP in a total volume of 100 μL. Samples were incubated at room temperature for 30 minutes, protected from light. Fluorescence was measured with a microplate reader (BMG LabTech); excitation 530–560 nm and emission detection at 590 nm.

### Mitochondrial membrane potential assay (JC-1)

Assays were performed using the JC-1 - Mitochondrial Membrane Potential Assay Kit (Abcam, ab113850, Cambridge, United Kingdom) according to the manufacturer’s instructions. After 24 h of pre-incubation, cells were treated with 0 or 100 ng/ml LPS for 3 h followed by nigericin (2μM) for 1 hour. Cells were washed once in Dilution Buffer and then stained with 20 μM JC-1 in 1X Dilution Buffer for 10 minutes at 37ºC. Cells were washed twice in 1X Dilution Buffer and fluorescence measured (Excitation 570 nm / Emission 620 nm).

### Nuclear translocation of p65

Peritoneal macrophages were seeded on glass coverslips in six well plates and treated with LPS for 3 h. Cells were washed three times with PBS and fixed with 4% formaldehyde in PBS for 10 min. After washing with PBS, the cells were permeabilized with 0.1% Triton X-100 in PBS for 1.5 min and incubated with blocking solution (3% BSA in PBS) for 1 h followed by incubating with the anti-p65 antibody for overnight. Cells were washed three times with PBS and incubated with anti-rabbit secondary antibody (Alexa Fluor 488; Molecular Probes) for 1 h in PBS in the dark. After washing three times with PBS, samples were mounted using antifade Gold reagent with DAPI (Invitrogen, P36935,Waltham, MA, USA). Immunofluorescence was detected by confocal microscopy. Co-localization of p65 and DAPI was quantified by ImageJ software. p65 translocation was quantified by the ratio of fluorescence pixel intensity in the nucleus (co-localization with DAPI) over the total cellular pixel intensity. At least 6 images were quantified for each experimental group (50).

### Cytokine assays

IL-6, MCP-1, IL-1 β and IL-18 in plasma and peritoneal fluids were determined by ELISA according to the respective manufacturer’s instructions. For in vitro studies, culture media was cleared by centrifugation at 16,000 ×g for 5 min and stored at −20 °C.

### Statistics

Unless otherwise noted, in vitro experiments were repeated as three independent procedures, with duplicate or triplicate wells averaged prior to statistical analysis. All data were presented as mean ± SEM. GraphPad Prism 9.0 (GraphPad Software Inc., San Diego, CA, USA) was used for statistical analysis. Comparisons between two groups after stimulation were analyzed by two-way ANOVA. Some experiments in cell cultures were analyzed by one-way ANOVA followed by post hoc T tests using Bonferroni correction for multiple comparisons. P values were indicated as follow: * < 0.05, ** < 0.01, *** < 0.001, **** < 0.0001.

## Author Contribution Statement

C.G.H., W.J.L., M.S., and B.C.B. designed research; C.G.H., W.J.L., C.L.C., and M.S. performed research; B.C.B. contributed new reagents/ analytic tools; C.G.H., and W.J.L. analyzed data; C.G.H., W.J.L., and B.C.B. wrote the paper.

## Acknowledgments

We thank Amanda Pereira and Sharon Senchanthisai for assistance with maintenance and breeding of mice.

## Ethics Statement

All of the experiments were approved by the University Committee on Animal Use For Research (UCAR) at the University of Rochester and followed National Institutes of Health guidelines for experimental procedures on mice.

## Data Availability Statement

All data generated or analysed during this study are included in this published article and in its supplementary file.

