## supplementary materials for "Pnpt1 mediates NLRP3 inflammasome activation by MAVS and metabolic reprogramming in macrophages"

Running title: Pnpt1 mediates inflammation in macrophages

Chia George Hsu[1,#] , Wenjia Li[1,2,#], Mark Sowden[1], Camila Lage Chávez[1], and Bradford C. Berk[1]

[1] Department of Medicine, Aab Cardiovascular Research Institute, University of Rochester, Rochester, NY, USA

[2]Department of Cardiology, Zhongshan Hospital, Fudan University, Shanghai, China

[#] These authors contributed equally to this work

Correspondence to

Bradford C. Berk, M.D., Ph.D.,

University of Rochester Medical Center

Box URNI, 601 Elmwood Ave

Rochester, NY 14642.

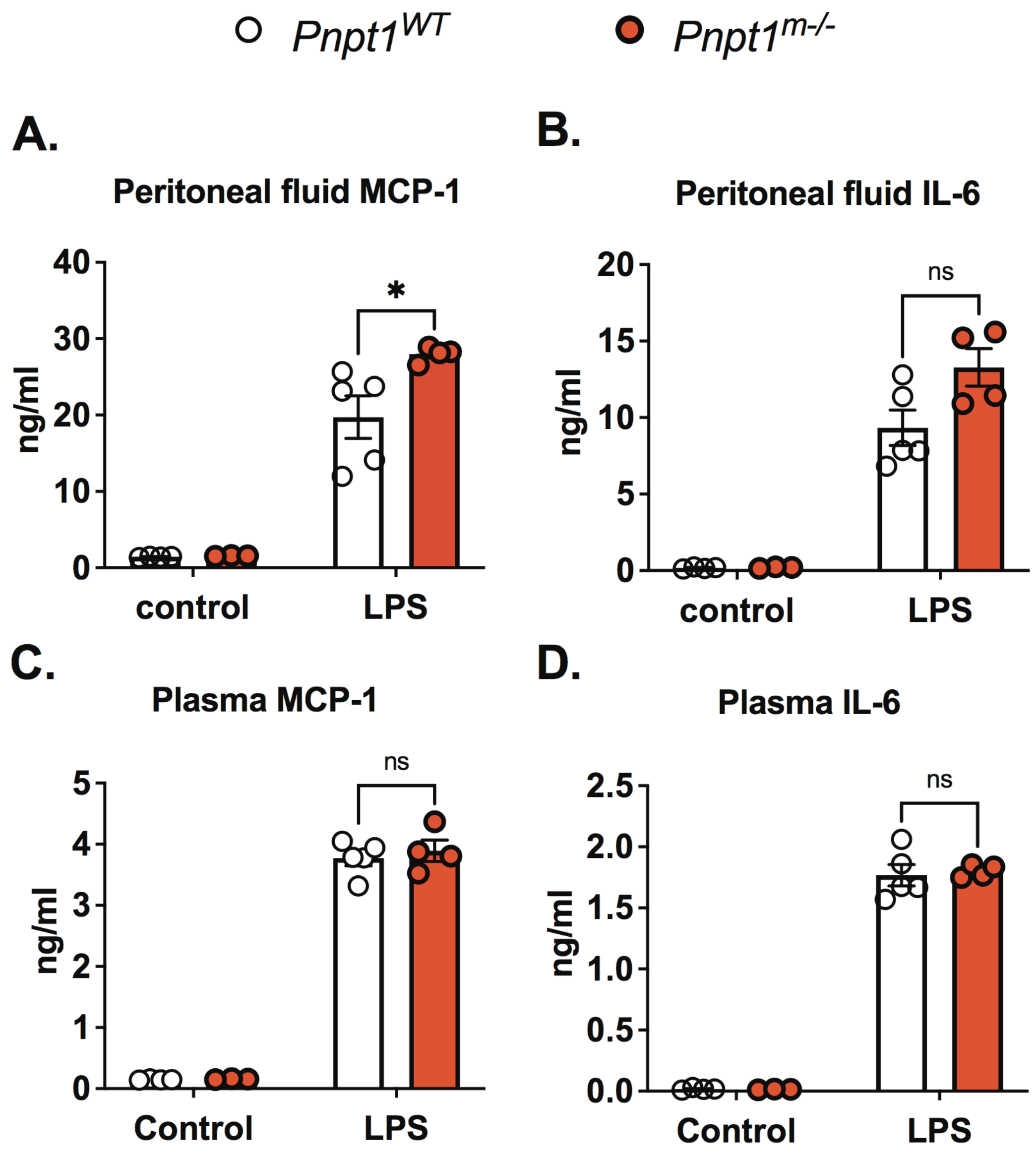


**Supplementary Fig. 1 Pnpt1 deletion has minimal effects on IL-6 and MCP-1 production**. Ten-week-old male Pnpt1$^{m-/-}$ and WT mice were sacrificed 6 h after i.p. injection with 40 mg/kg of LPS. The whole blood samples were collected and peritoneal cavities were washed with PBS. (A) MCP-1 and (B) IL-6 in peritoneal lavage fluid; (C) MCP-1 and (D) IL-6 in plasma (n=4-5 mice per group). Statistics in B-E were performed using a 2-way ANOVA and Bonferroni's post hoc test. *P<0.05 between LPS-WT and LPS-Pnpt1$^{m-/-}$ groups. Bars represent mean ± SEM. Statistics in B-C were performed using a two-way ANOVA and Bonferroni's post hoc test. Bars represent mean ± SEM.   *p <0.05.

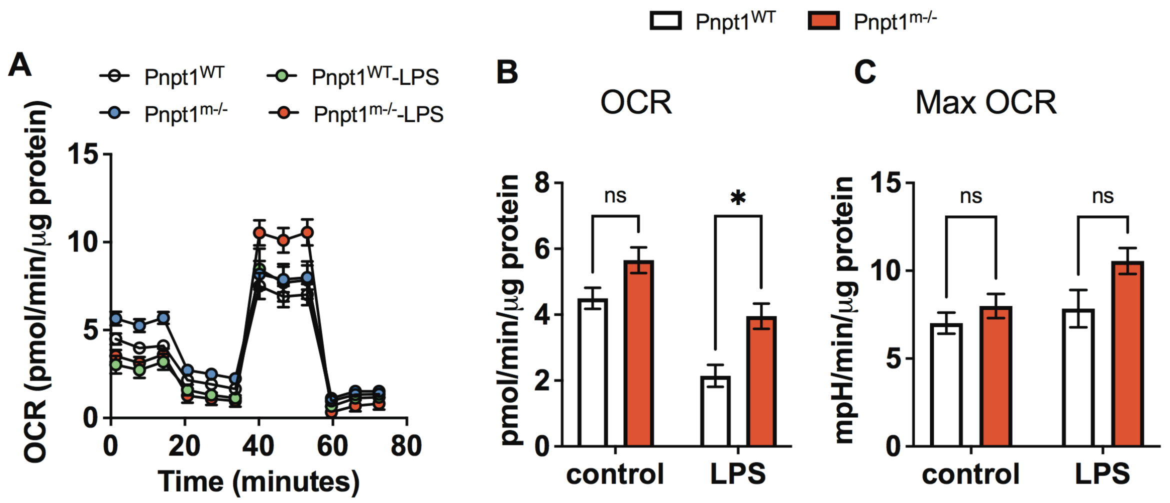


**Supplementary Fig. 2. Pnpt1 deletion increases oxygen consumption.** Untreated (Con) or LPS-activated (3 hr) peritoneal macrophages from Pnpt1$^{m-/-}$ and WT mice were subjected to a mitochondrial stress test using a Seahorse XF-analyzer. (A) A representative seahorse plot of the mitochondrial stress test assessed by oxygen consumption rate (OCR) , an index of glycolysis after injection of oligomycin (1 µg/ml), carbonyl cyanide-4-(trifluoromethoxy)phenylhydrazone (FCCP, 1 µM), and rotenone (1 µM) plus antimycin (1 µM). (B) Basal OCR (C) Maximal OCR. Bars represent mean ± SEM. Statistics in B-C were performed using a two-way ANOVA and Bonferroni's post hoc test. Bars represent mean ± SEM. *$p < 0.05$.

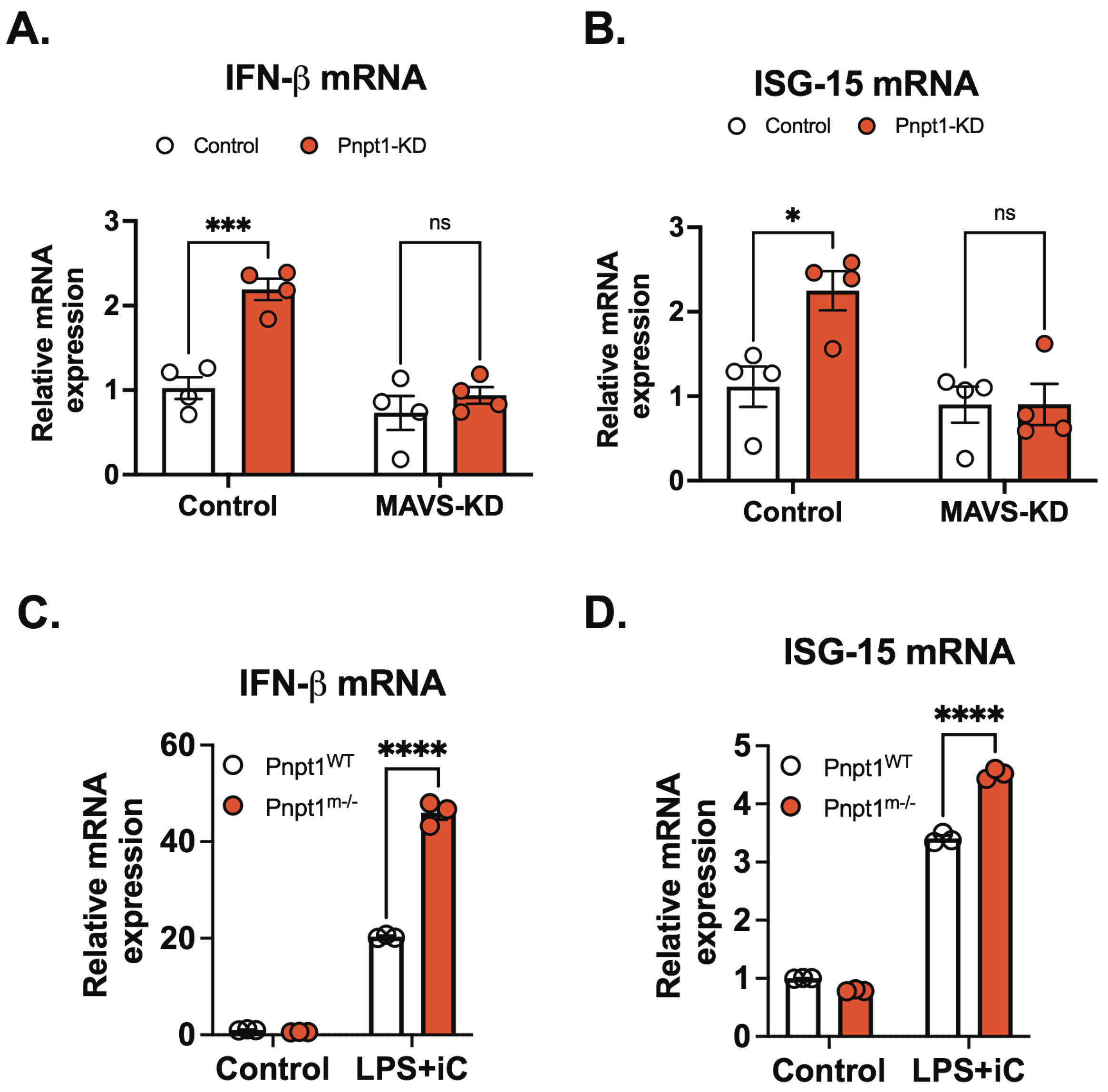


**Supplementary Fig. 3. Pnpt1 deficiency enhances Type-I Interferon Responses.** HEK-293T cells were knocked down with scramble (control), Pnpt1 or MAVS siRNA for 48 hr., and gene expression was measured by real-time PCR. (A) IFN-β (B) ISG-15 mRNA from cell lysates (n=4 experiments). BMDMs from Pnpt1m-/- and WT mice were stimulated with LPS (100 ng/mL) for 3 hr. followed by 10 μM Poly(I:C) transfection (iC) for 12 hr. (A) IFN-β (B) ISG-15 mRNA from cell lysates (n=3 experiments). (Statistics in A-D were performed using a 2-way ANOVA and Bonferroni's post hoc test. Bars represent mean ± SEM. *p <0.05, ***p <0.001, ****p <0.001.
